# Activating FcγRs on monocytes are necessary for optimal Mayaro virus clearance

**DOI:** 10.1101/2024.07.23.604823

**Authors:** Megan M. Dunagan, Nathânia Dábilla, Colton McNinch, Jason M. Brenchley, Patrick T. Dolan, Julie M. Fox

## Abstract

Mayaro virus (MAYV) is an emerging arbovirus. Previous studies have shown antibody Fc effector functions are critical for optimal monoclonal antibody-mediated protection against alphaviruses; however, the requirement of Fc gamma receptors (FcγRs) for protection during natural infection has not been evaluated. Here, we showed mice lacking activating FcγRs (FcRγ^-/-^) developed prolonged clinical disease with more virus in joint-associated tissues. Viral clearance was associated with anti-MAYV cell surface binding rather than neutralizing antibodies. Lack of Fc-FcγR engagement increased the number of monocytes through chronic timepoints. Single cell RNA sequencing showed elevated levels of pro-inflammatory monocytes in joint-associated tissue with increased MAYV RNA present in FcRγ^-/-^ monocytes and macrophages. Transfer of FcRγ^-/-^ monocytes into wild type animals was sufficient to increase virus in joint-associated tissue. Overall, this study suggests that engagement of antibody Fc with activating FcγRs promotes protective responses during MAYV infection and prevents monocytes from being potential targets of infection.

## Introduction

Alphaviruses are transmitted by mosquitoes and have caused explosive outbreaks worldwide [1]. Arthritogenic alphaviruses, including chikungunya virus (CHIKV), Ross River virus (RRV), and Mayaro virus (MAYV), can cause fever, myalgia, and arthralgia. Up to 50% of infected individuals can develop polyarthralgia lasting months to years following initial infection, with rare cases showing neurological complications or even death [2–5]. Since its identification in 1954, MAYV has caused occasional outbreaks in rural areas of Central and South America as well as the Caribbean [6]. While MAYV is primarily transmitted by forest dwelling *Haemagogus sp.* mosquitoes, *Aedes aegypti* have been shown experimentally to be competent vectors for MAYV, highlighting the potential for MAYV to spread into more populated urban regions [7]. Despite these risks MAYV remains understudied, as cases are largely underreported or misdiagnosed [8–10]. Understanding the aspects of immunity that contribute to pathogenesis and disease resolution will help inform the development of vaccines and therapeutics.

Alphaviruses are enveloped, positive-sense RNA viruses with a genome encoding four nonstructural proteins (nsP1-nsP4) and six structural proteins (capsid, E3, E2, 6K, TF, and E1) [11]. Following viral replication, trimers of p62 (E3 and E2) and E1 are assembled and trafficked to the cell surface for subsequent budding of the progeny virions [12]. During transport through the trans-Golgi network, furin-like proteases cleave E3 to produce the mature E2-E1 heterodimer [13]. The E2 and E1 surface glycoproteins mediate viral attachment and fusion, respectively, and have been characterized broadly as targets of the antibody responses following alphavirus infection [14–19]. Anti-alphavirus antibodies have been shown to block multiple stages in the viral life cycle including attachment, entry, fusion, and egress [18, 20, 21]. Furthermore, anti-alphavirus antibodies can bind to the E2 and E1 proteins present on the infected cell surface, in addition to free virions, and mediate enhanced clearance and immune modulation through Fc interaction with host proteins [*e.g*., Fc gamma receptors (FcγRs) and the complement component, C1q] [22–24].

Mouse models of alphavirus disease recapitulate aspects of human infection. MAYV infection in an immunocompetent [C57BL/6 wild type (WT)] mouse model results in high viral titers, symmetric joint swelling, and a robust innate and adaptive immune response [19, 25]. Similar to mouse models of CHIKV and RRV, antibodies are necessary to clear circulating infectious MAYV; T and B cell deficient (RAG^-/-^) mice survive MAYV infection with sustained viremia, and administration of cross-reactive alphavirus immune serum suppresses MAYV viremia to undetectable levels for a short duration [26–28]. Previous studies evaluating monoclonal antibody (mAb) efficacy against CHIKV or MAYV showed a requirement for Fc-mediated activity for optimal protection [18, 19, 23, 29]. For MAYV, the necessity of Fc effector functions for protection was independent of time of mAb administration and *in vitro* neutralization potency [19, 30]. These studies clearly highlight the importance of Fc-FcγR interactions for mAb-mediated protection during alphavirus infection. Of note, these studies administered mAbs either before or within a few days of infection, which is prior to the generation of an endogenous humoral response. As such, the contribution of Fc-FcγR interactions during primary MAYV infection remains unclear.

Here, we evaluated the role of Fc-FcγR interactions for disease resolution during a primary MAYV infection using mice that lack the Fc common gamma chain (FcRγ^−/−^) and thus do not express activating FcγRs [31]. Mice lacking activating FcγRs showed prolonged foot swelling and increased viral RNA and infectious virus during disease resolution, despite having similar levels of binding and neutralizing antibodies. Infection of B cell-depleted or mice lacking mature B cells demonstrated the necessity of FcγR interaction with anti-MAYV reactive antibodies for MAYV clearance, rather than neutralizing antibodies. FcRγ^−/−^ mice had increased infiltration of immune cells into the joint-associated tissue during acute disease with an altered proportion of monocytes to macrophages which persisted to a chronic time point. Analysis of the myeloid cell populations by single cell RNA sequencing (scRNAseq) showed increased viral RNA in monocytes and macrophage clusters, which corresponded with enrichment of pathways associated with type I IFN signaling, antiviral response, and cellular stress response. Adoptive transfer of FcRγ^−/−^ monocytes was sufficient to increase viral burden in WT mice. Overall, these studies indicate that Fc-FcγR interactions are necessary for optimal MAYV clearance and disease resolution and FcγR engagement on monocytes may impact the susceptibility of these cells to MAYV infection.

## Results

### Activating FcγRs enhance disease resolution and viral clearance during MAYV infection

In the immunocompetent mouse model of MAYV-induced musculoskeletal disease, MAYV inoculation in footpad results in swelling of both the infected (ipsilateral) and contralateral foot through 8 days post-infection (dpi), with peaked planar edema between 5 to 6 dpi in the ipsilateral foot and infectious virus measurable through 10 dpi [25]. To assess the contribution of activating FcγRs during MAYV infection, we inoculated four-week-old C57BL/6N WT or FcRγ^−/−^ mice subcutaneously in the rear footpad with 10^3^ focus forming units (FFU) of MAYV and measured foot swelling through 25 dpi. As expected, swelling of the ipsilateral and contralateral feet peaked between 5 to 6 dpi and substantially decreased by 8 dpi in WT mice (**Fig. 1a** and **Extended Data Fig. 1a**). While foot swelling still peaked at 5 to 6 dpi in FcRγ^-/-^ mice, with no difference in the overall magnitude of swelling, the absence of activating FcγRs lead to prolonged swelling until 17 dpi in the ipsilateral foot (**Fig. 1a**) and 11 dpi in the contralateral foot (**Extended Data Fig. 1a**).

**Figure 1.**
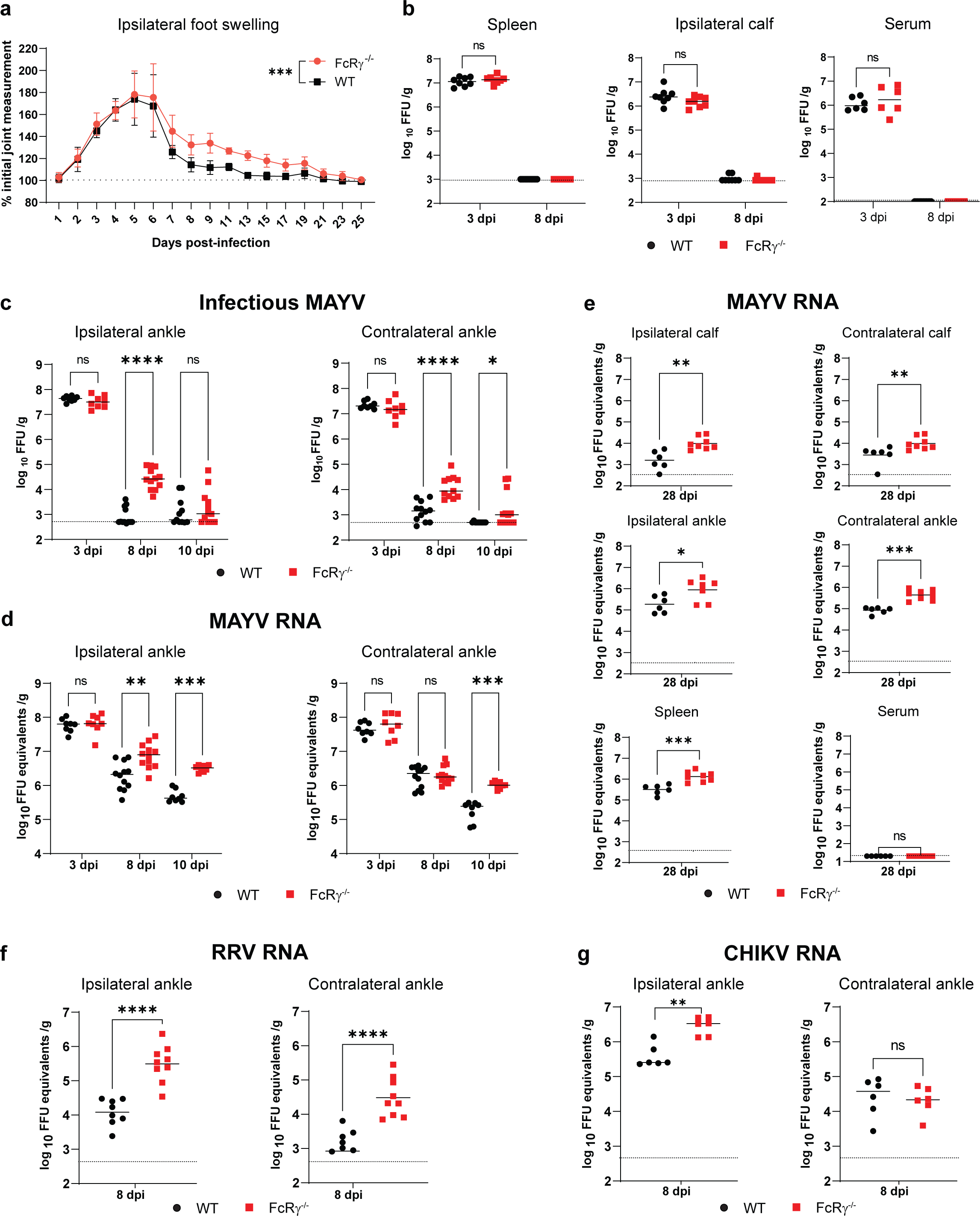
Prolonged foot swelling and viral burden in joint-associated tissue of FcRγ^-/-^ mice following MAYV infection. Four-week-old WT or FcRγ^−/−^ C57BL/6N mice were infected subcutaneous in the rear footpad with 10^3^ focus forming units (FFU) of (**a-e**) MAYV, (**f**) RRV, or (**g**) CHIKV. (**a**) Swelling of the ipsilateral foot was measured prior to infection and for 25 dpi (n = 8 per group, 2 independent experiments). Graphs show mean ± SEM. Statistical significance was determined based on area under the curve (AUC) analysis using student’s t-test ***, *P* < 0.001. (**b-g**) Indicated tissues were harvested at 3, 8, or 10 dpi and titrated for (**b, c**) infectious virus by focus forming assay (FFA) or (**d-g**) viral RNA by qRT-PCR with virus specific primers and probe (n = 8 to 12 per group; 2 to 3 independent experiments). Statistical significance was determined by a Mann-Whitney test. (*, *P* <0.05; **, *P* < 0.01; ***, *P* < 0.001; ****, *P* < 0.0001; ns = not significant.) Bars indicate the median value and dotted lines indicate the limit of detection for the assay.

Following infection, MAYV rapidly disseminates causing high viremia and viral burden in skeletal muscles, spleen, and joint-associated tissues by 3 dpi. Similar levels of infectious virus were observed at 3 dpi in spleen, gastrocnemius (calf) muscle, and serum of WT and FcRγ^−/−^ mice (**Fig. 1b**). By 8 dpi, infectious virus was not detectable in these tissues but there was a significant increase in viral RNA in the spleens of FcRγ^−/−^ mice (**Extended Data Fig. 1b**). We next quantified infectious MAYV and MAYV RNA from joint-associated tissues. While there were similar viral loads at 3 dpi, FcRγ^−/−^ mice showed delayed clearance of infectious virus and viral RNA in the ipsilateral and contralateral ankles at 8 and 10 dpi compared to WT mice (**Fig. 1c and d**). The delay in viral clearance was maintained at a chronic time point (28 dpi), with FcRγ^−/−^ mice having significantly more MAYV RNA in the ankles, calves, and spleen compared to WT mice (**Fig. 1e**). These data suggest that clearance defects resulting from the lack of activating FcγRs are most notable the joint-associated tissue during acute infection but can also persist at the RNA level in other tissues during chronic time points.

To determine if FcγR-mediated viral clearance applied more broadly to arthritogenic alphaviruses, we infected WT or FcRγ^-/-^ mice with either RRV or CHIKV and quantified viral RNA from the ankles at 8 dpi (**Fig. 1f and g**). Viral RNA levels were increased in FcRγ^−/−^ mice compared to WT mice following RRV and CHIKV infection suggesting that activating FcγRs are more broadly required for optimal clearance of viral RNA during primary alphavirus infection.

### FcγR interaction with cell surface binding antibodies is necessary for MAYV clearance

Previous studies have highlighted the importance of antibodies for the clearance of infectious alphaviruses [18, 30, 32]. Despite the lack of activating FcγRs on B cells, variations in the tissue microenvironment could impact the anti-MAYV antibody response following infection in the FcRγ^−/−^ mice [31, 33]. To determine if the delay in MAYV clearance was related to altered antibody titers or neutralization potency, we quantified MAYV E2-specific IgG by ELISA and neutralizing antibodies by FRNT at 3, 8, 10, and 28 dpi (**Fig. 2a**). As expected, no anti-MAYV antibodies were detectable in circulation at 3 dpi. At 8 dpi, FcRγ^-/-^ mice had significantly lower levels of neutralizing antibodies, but equivalent antibody neutralization titers between the groups by 10 dpi. Despite the early difference in neutralizing antibodies, there was no difference in either total anti-E2 IgG or neutralizing antibodies detected by 28 dpi (**Fig. 2a**). These results indicate that, while there is a delay in the generation of specifically neutralizing antibody, there is no dramatic defect in antibody response in FcRγ^-/-^ mice.

**Figure 2:**
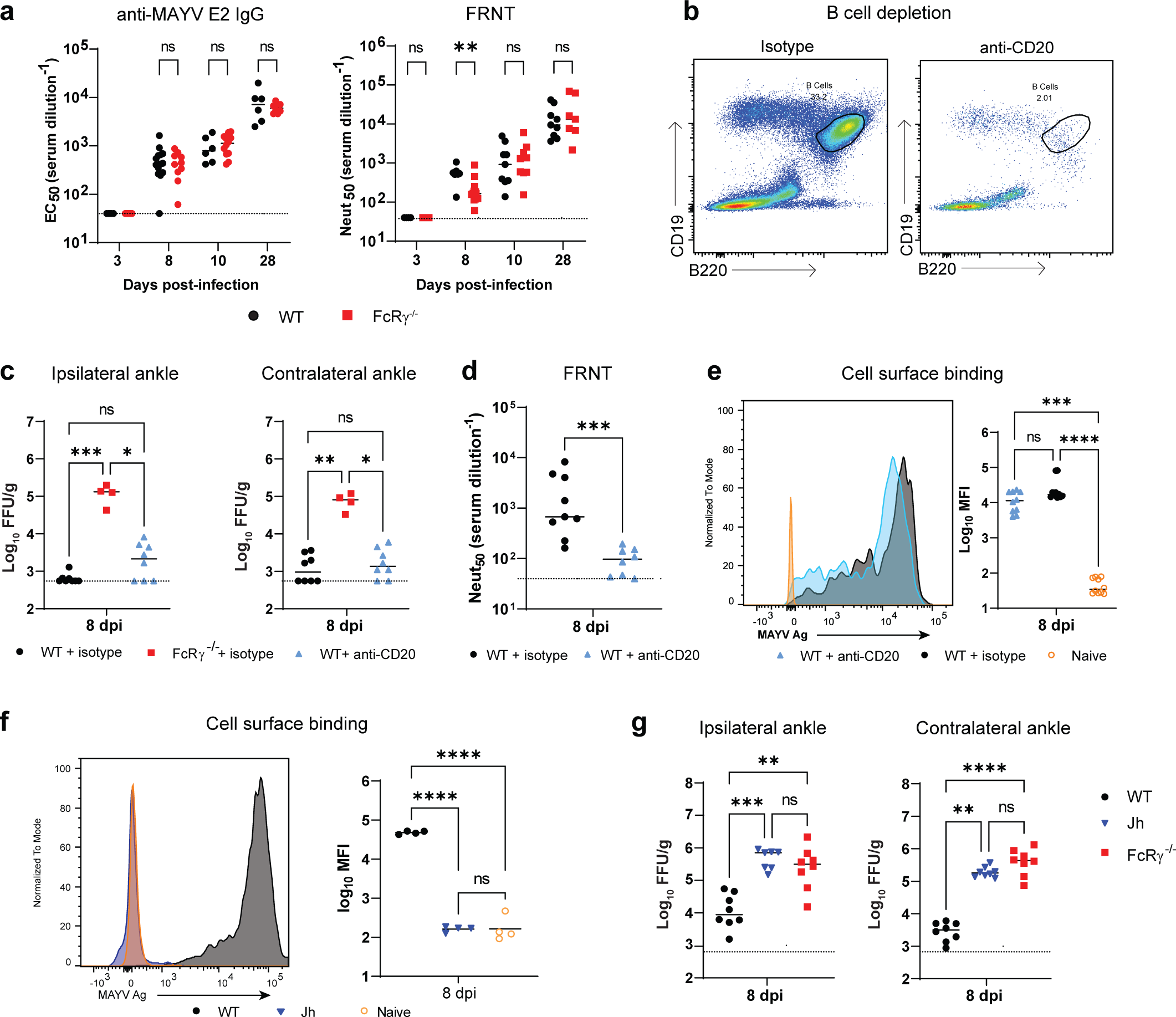
Interaction of cell surface binding anti-MAYV antibodies with activating FcγRs mediate MAYV clearance in joint-associated tissues. Four-week-old WT or FcRγ^−/−^ C57BL/6N mice were infected subcutaneous in the rear footpad with 10^3^ FFU of MAYV. (**a**) Serum was collected at indicated time points. Serial dilutions of serum were used to determined EC_50_ values of anti-MAYV E2-specific IgG by ELISA and Neut_50_ values for MAYV neutralization by focus reduction neutralization test (FRNT) (n = 8 to 15 per group; 2 to 3 independent experiments). (**b-e**) B cells were depleted using anti-CD20 administered on -1 and 4 dpi (500 µg/dose) and control mice received a non-depleting isotype control antibody (n = 4 to 8 per group; 2 independent experiments). (**b**) PBMCs were collected at 8 dpi to confirm B cell (CD19^+^B220^+^) depletion by flow cytometry. (**c**) Infectious virus was titrated from the ipsilateral and contralateral ankles by FFA at 8 dpi. (**d**) Neut_50_ values of serum tested for MAYV neutralization by FRNT. (**e**) Serum antibodies binding to the surface of live MAYV-infected Vero cells was evaluated by flow cytometry. (**f-g**) Jh, WT, or FcRγ^-/-^ mice were infected with MAYV and tissues were collected at 8 dpi. (**f**) Cell surface MAYV-binding antibodies were quantified from the serum and (**g**) infectious virus was titrated from the ankles (n = 4 to 8; 2 independent experiments). Bars indicate the median values. Statistical significance was determined by a Mann-Whitney test (a and d) or one-way ANOVA (c, e, f, g). *, *P* <0.05; **, *P* < 0.01; ***, *P* < 0.001; ****, *P* < 0.0001; ns = not significant.

To confirm the requirement of antibodies for the delayed viral clearance in the FcRγ^-/-^ mice, we depleted B cells using an anti-CD20 antibody, as previously described [34]. Since the mechanism of anti-CD20 is dependent on activating FcγRs, we could not deplete B cells in the FcRγ^−/−^ mice. Instead, WT mice administered an anti-CD20 antibody or isotype control were compared to FcRγ^−/−^ mice administered an isotype control. Consistent with previous data, anti-CD20 antibody treatment dramatically decreased double positive B220^+^CD19^+^ B cells from the blood of infected mice at 8 dpi (**Fig. 2b**). Surprisingly, B cell depletion failed to increase infectious virus burden to the level observed in FcRγ^−/−^ mice. Instead, viral burden recapitulated WT mice administered an isotype control (**Fig. 2c**). As a secondary measure of B cell depletion, we quantified levels of neutralizing and cell surface binding antibodies in the serum at 8 dpi. While B cell depletion significantly reduced the level of neutralizing antibodies (**Fig. 2d**), there was no difference in the amount of IgG in the serum that could bind to the surface of MAYV-infected Vero cells compared to the isotype control (**Fig. 2e**). These results show there was still significant anti-MAYV antibody generated despite anti-CD20 depletion, which is most likely due to a combination of incomplete B cell depletion from tissues and recovery of B cell populations following depletion [35]. However, the substantial reduction in neutralizing antibodies only marginally impacted the clearance of infectious virus from joint-associated tissue suggesting a more dominant role for antibodies that bind the surface of infected cells regardless of neutralizing ability.

To fully address the contribution of soluble antibody to MAYV clearance, we infected Jh (C57BL/6N) mice with MAYV and quantified infectious virus in the ankles at 8 dpi. Jh mice lack the J segment of the Ig heavy chain, a mutation that stalls B cell development at the precursor stage [36] and thus cannot produce antibodies. Indeed, serum collected from Jh mice at 8 dpi did not contain IgG that bound to the surface of MAYV-infected Vero cells (**Fig. 2f**). Jh mice failed to efficiently clear infectious virus from the ankles as compared to the WT mice (**Fig. 2g**). Interestingly, the viral burden in ankles of Jh mice was similar to the viral load in FcRγ^−/−^ mice even though FcRγ^−/−^ mice still have neutralizing antibodies present (**Fig. 2a and g**). Taken together, these results suggest that MAYV clearance from joint-associated tissues depend more on the interaction of activating FcγRs with, presumably, the Fc region of antibodies bound to the surface of infected cells rather than the presence of neutralizing antibodies.

### Prolonged monocyte recruitment to the site of infection in the absence of activating FcγRs

Recruitment of inflammatory immune cells has been implicated in both protection as well as immunopathology following alphavirus infection. While little is known about cellular contributors to disease during MAYV infection, infiltrating CD4^+^ T cells and monocytes/macrophage have been shown to contribute to disease in CHIKV models and FcγR engagement with anti-CHIKV mAbs was shown to alter immune cell infiltration [22, 37]. To determine if FcγR engagement with the endogenous humoral response alters the flux of immune cells, the ipsilateral foot was harvested from MAYV-infected WT or FcRγ^-/-^ mice at 3, 8, 10, or 28 dpi. Following digestion, single cell suspensions were stained and analyzed by flow cytometry (**Extended Data Fig. 2a**). In mice, activating FcγRs are expressed predominantly on monocytes, macrophage, DCs, and various granulocytes, leading us to focus our analysis on these cellular subsets [33]. There was a similar number of CD45^+^ cells between the groups (**Extended Data Fig. 2b**). However, mice lacking activating FcγRs had increased numbers of neutrophils, NK cells, and Ly6c^hi^ monocytes between 8-10 dpi (**Fig. 3a**). FcRγ^−/−^ mice also had increased numbers of CD8^+^ T cells and B cells, but not CD4^+^ T cells. (**Extended Data Fig. 2b and c**). In other alphavirus mouse models, CD8^+^ T cells do not contribute to viral clearance or disease resolution specifically in joint-associated tissue, so we did not investigate this response further [38, 39]. Interestingly, WT mice had proportionally more Ly6c^mid-low^F4/80^+^ macrophages and fewer Ly6C^hi^ monocytes compared to FcRγ^−/−^ mice at 10 dpi (**Fig. 3b**). While most immune cell populations had returned to within naïve ranges at 28 dpi, there were significantly more monocytes with a corresponding reduction in macrophage in FcRγ^−/−^ mice compared to WT mice (**Fig. 3c**). The proportion of macrophages in the FcRγ^−/−^ tissue was even below naïve levels suggesting a defect in the return of cellular homeostasis in the absence of activating FcγRs (**Fig. 3d**). Taken together, these results show that the lack of activating FcγRs impacts the monocytes and macrophages dynamics during infection and recovery.

**Figure 3.**
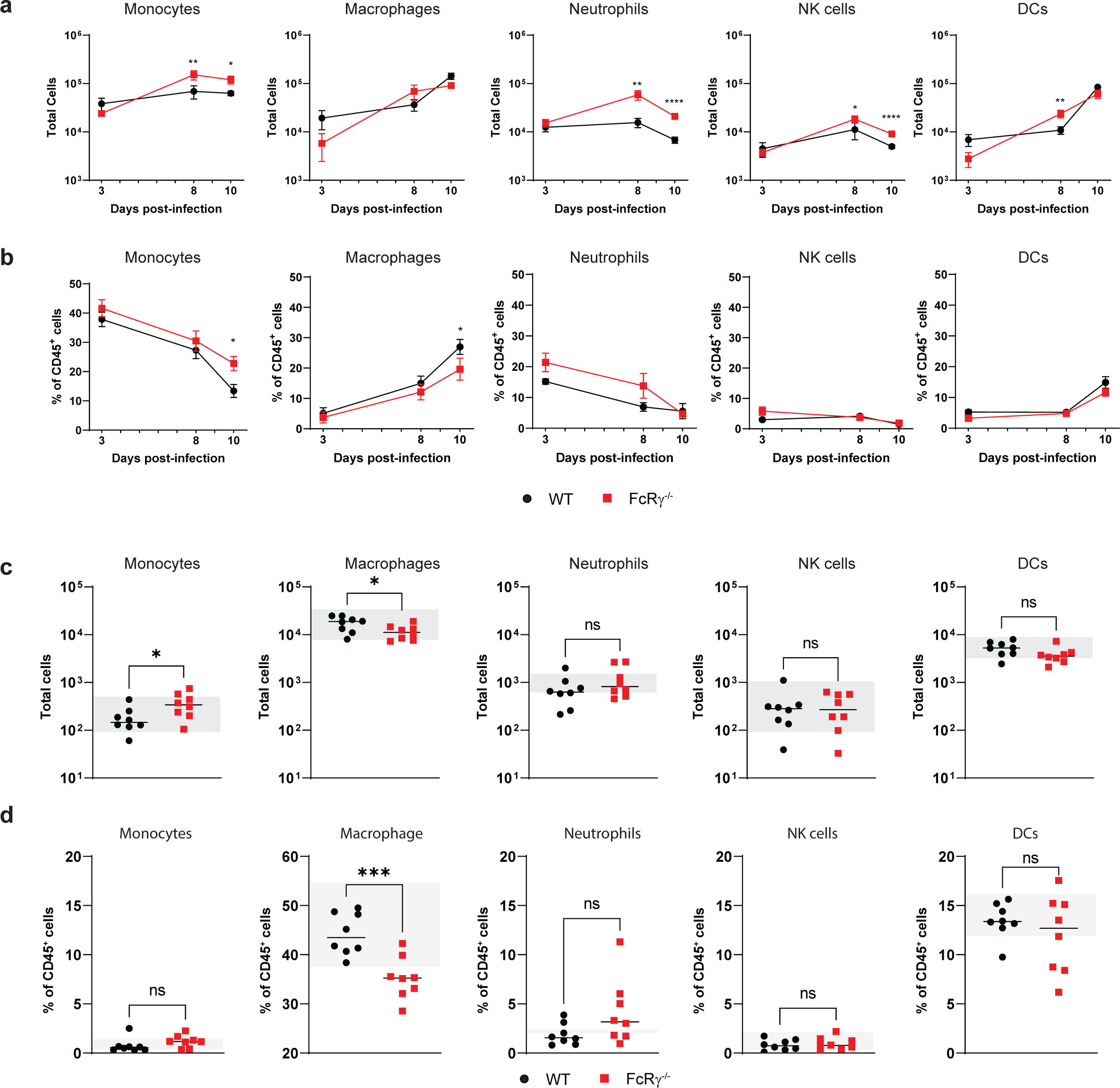
Lack of activating FcγRs alters flux of immune cells in the ipsilateral foot. Four-week-old WT or FcRγ^−/−^ C57BL/6N mice were infected subcutaneous in the rear footpad with 10^3^ FFU of MAYV. Single cell suspensions were isolated from the ipsilateral foot and proximal skin at (**a-b**) 3, 8, and 10 dpi or (**c-d**) 28 dpi and stained for monocytes (Ly6C^hi^), macrophages (Ly6C^mid-lo^F4/80^+^), neutrophils (Ly6G^+^), NK cells (NK1.1^+^), and dendritic cells (DCs; CD11b^-^ CD11c^+^ MHCII^+^) and analyzed by flow cytometry to determine the (**a, c**) total numbers of viable cells or (**b, d**) percentage of CD45^+^ cells (n = 5 to 8 per group; 3 independent experiments). (**c, d**) The gray bar represents the range of (**c**) total cells and (**d**) percentage of CD45^+^ cells from WT and FcRγ^−/−^ naïve mice. Graphs show mean ± SEM. Statistical significance was determined using a Mann-Whitney test at individual time points test. *, *P* <0.05; **, *P* < 0.01; ***, *P* < 0.001; ****, *P* < 0.0001; ns = not significant.

### Increased MAYV RNA in monocyte and macrophages without activating Fc**γ**Rs

The shift in the monocyte and macrophage populations in the FcRγ^-/-^ mice suggests there might be a differential transcriptional profile between the cellular populations. Additionally, previous studies have implicated monocytes and macrophages as potential targets of alphavirus infection which may impact the cellular response [40–42]. To evaluate alterations in immune cell profiles and the presence of MAYV in specific immune cells, we performed single cell RNA sequencing on the ipsilateral foot from naïve or MAYV-infected WT or FcRγ^−/−^ mice at 10 dpi. Single cell suspensions were stained, and leukocytes were sorted based on CD45 expression. CD45^+^ cells were subjected to micro-fluidics-based single-cell RNA sequencing with the addition of MAYV-specific primers. More than 2,000 cells were collected from each group and > 100,000 RNA reads were analyzed per cell. Immune cell subsets were identified based on key markers and expert curation (**Extended Data Fig. 3a-c**) [43–67]. Integrated analysis of all samples showed the presence of distinct immune cell clusters encompassing similar cellular populations identified in our flow cytometry analysis (**Fig. 4a**). These populations included NK cells, B cells, CD8^+^ and CD4^+^ T cells, macrophages, monocytes, neutrophils, and dendritic cells with the addition of mast cells and γδ T cells. The proportion and enrichment of each cluster was consistent between replicates from each experimental group highlighting the reproducibility of the analysis (**Extended Data Fig. 3d**). We next evaluated the distribution of MAYV RNA within the integrated data set. Individual cells expressing MAYV RNA are colored as per their cluster and the size of the cell point indicates to the level of viral RNA. There was a clear enrichment of viral RNA in myeloid cell clusters including Clusters 0, 1, 2, 3, 4, and 8 (**Fig. 4b**). Cluster enrichment analysis showed an increased proportion of cells in Clusters 0-2 contained viral RNA (**Fig. 4c**).

**Figure 4.**
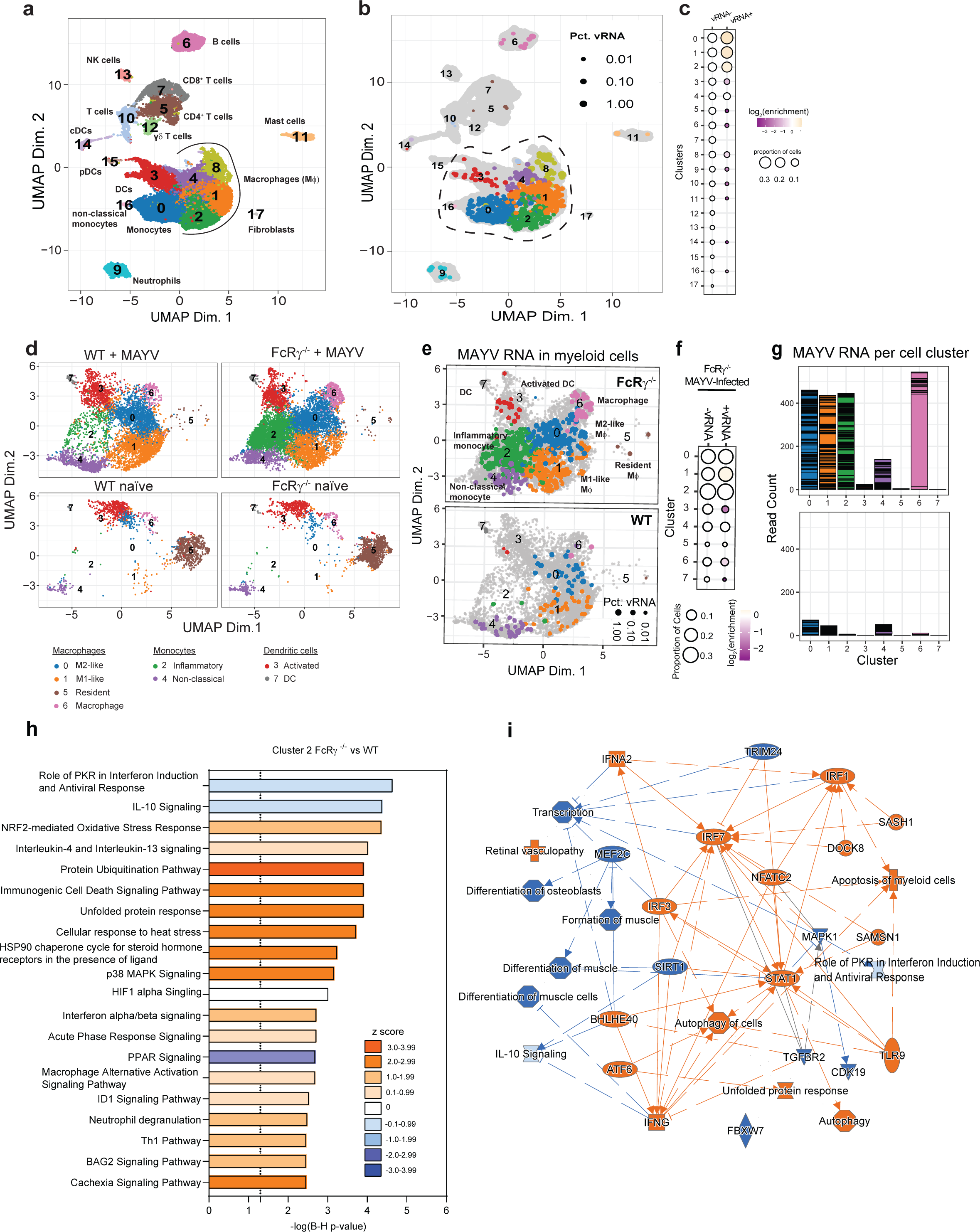
Increased MAYV RNA in myeloid cells lacking activating FcγRs. Four-week-old WT or FcRγ^−/−^ C57BL/6N were infected subcutaneous in the rear footpad with 10^3^ FFU of MAYV or mock infected with diluent. At 10 dpi, the ipsilateral foot and surrounding skin was enzymatically digested into a single cell suspension and stained for CD45. Viable CD45^+^ cells were sorted then subjected to microfluidic-based single cell RNA sequencing with the addition of MAYV-specific primers. Over 2000 cells were collected per group with >100,000 RNA reads per cell. (**a**) UMAP shows the integrated cell clusters from all groups. (**b**) Cells containing MAYV RNA are identified based on the cluster color in (a). The size of the dot represents the percentage of viral RNA in the cells. (**c**) The proportion of cells across each integrated cluster segregated based on the presence of MAYV RNA indicating the log_2_(enrichment) of each cluster between viral RNA positive and negative cells. (**d**) Subcluster analysis was performed on the myeloid cell subset, which is indicated by the dotted line in (b). (**e**) Cells expressing MAYV RNA in the myeloid subcluster are colored based on the subcluster analysis and the size denotes the percentage of viral RNA in the cell. (**f**) The proportion of cells from the FcRγ^-/-^ myeloid subclusters separated by the presence of MAYV RNA showing the log_2_(enrichment) for each cluster between viral RNA positive and negative cells. (**g**) Total MAYV RNA read count per cell in the myeloid subcluster analysis. Each horizontal bar represents a single cell. (**h-i**) Ingenuity pathway analysis on differentially expressed genes in FcRγ^-/-^ monocytes (Cluster 2) using an enrichment cutoff of Log2FC ≤ -0.58 or ≥ 0.58 and an adjusted P value < 0.05. (**h**) Enriched canonical pathways and (**i**) graphical summary of predicted pathway activity in FcRγ^-/-^ monocytes compared to WT mice. (**h**) Orange bars indicate a positive z-score, blue bars indicate a negative z-score, and white bars represent a z-score of 0. (**i**) Orange nodes/lines indicate predicted activation and blue nodes/lines indicate predicted inhibition. Relationships between nodes are distinguished by the lines. A solid line leads to activation/inhibition, a thin dashed line is an inferred relationship, and a grey line represents a direct interaction.

Focusing on the cellular populations with MAYV RNA, we reanalyzed the clusters outlined in the black dotted line (Clusters 0, 1, 2, 3, 4, 8, 16) (**Fig. 4b**) and identified the myeloid cell subsets using key gene markers (**Fig. 4d** and **Extended Data Fig. 4a-c**). A clear distinction between MAYV-infected and naïve mice was observed, with the loss of tissue-resident macrophages (Cluster 5) and an increase in the clusters identified as monocytes, activated macrophages, and dendritic cells (**Fig. 4d** and **Extended Data Fig. 4d**). There were no substantial differences in the clusters identified in the naïve groups, suggesting similar myeloid populations are present at baseline (**Extended Data Fig. 4d**). Compared to infected WT mice, infected FcRγ^-/-^ mice had a dramatic increase in the number of inflammatory monocytes (Cluster 2), which is consistent with our flow cytometry results (**Fig. 3a and b**). Evaluation of MAYV RNA distribution revealed an increased presence and enrichment of viral RNA in the FcRγ^-/-^ mice across multiple clusters (**Fig. 4e-g** and **Extended Data Fig. 4e**). M1-like macrophages (Cluster 1) were enriched in the viral RNA positive cells compared to the negative cells. This was distinct from the WT mice where non-classical monocytes (Cluster 4) were enriched in the viral RNA positive cells (**Extended Data Fig. 4e**). For both groups, there was a negative enrichment for Cluster 3 – 7 in the viral RNA positive cells suggesting these cells are actively preventing infection (**Extended Data Fig. 4e**). In WT mice, inflammatory monocytes (Cluster 2) were also negatively enriched in viral RNA containing cells; however, this may be due to the reduced number of cells present in the cluster (**Extended Data Fig. 4e**). On a per cell basis, the macrophages (Clusters 0, 1, and 6) and monocytes (Clusters 2 and 4) from FcRγ^-/-^ mice had the most MAYV RNA reads, albeit the majority of the MAYV reads in Cluster 6 are derived from one cell (**Fig. 4g**). Although the MAYV RNA read count was lower in the WT mice, most of the reads grouped within Clusters 0, 1, and 4 suggesting a differential enrichment of MAYV RNA in inflammatory monocytes (Clusters 2) in the FcRγ^-/-^ mice (**Fig. 4g**).

To interrogate the differences in the monocyte (Cluster 2) response, which may have contributed to the increased viral RNA, we performed ingenuity pathway analysis (IPA) on differentially expressed genes (DEGs) enriched in the FcRγ^-/-^ mice compared to WT mice. Up-regulated pathways (z score > 0) included broad categories of cellular stress response, protein ubiquitination pathways, type I interferon signaling, cytokine and chemokine signaling, and cellular differentiation (**Fig. 4h and i**). The down-regulated pathways (z score < 0) involved protein kinase R (PKR) induction, IL-10 signaling, and peroxisome proliferator activating receptor (PPAR) signaling (**Fig. 4h and i**). When enriched pathways were compared between the myeloid subclusters from the FcRγ^-/-^ mice, similar gene ontology pathways were identified, suggesting that the transcriptional landscape within these cell populations is similar (**Extended Data Fig. 5**). Overall, these results show enriched MAYV RNA in myeloid cells that lack activating FcγRs, which correlated with increased cellular stress and type I interferon response.

### Monocytes lacking activating Fc**γ**Rs are sufficient to drive prolonged MAYV infection

Earlier results showed that FcRγ^−/−^ mice had increased Ly6C^hi^ monocytes in the ipsilateral foot through 28 dpi (**Fig. 3**) and contained more MAYV RNA at 10 dpi (**Fig. 4**). Monocytes have previously been implicated in both tissue damage and/or disease resolution following alphavirus infection and have been shown to be productive targets of MAYV infection [37, 41, 68, 69]. To determine if FcRγ^−/−^ monocytes were sufficient to increase MAYV in the joint-associated tissue, FcRγ^−/−^ monocytes (CD45.2) were enriched from the bone marrow of donor mice, transferred into MAYV-infected CD45.1 (WT) or CD45.2 (FcRγ^−/−^) recipient mice at 0 and 4 dpi, and MAYV was quantified from the ipsilateral ankle of recipient mice at 8 dpi (**Fig. 5a**). The presence of transferred cells in WT animals was confirmed in the spleen and contralateral ankle at 8 dpi by flow cytometry (representative plot, **Fig. 5b**). Transfer of the FcRγ^−/−^ monocytes significantly increased the level of MAYV RNA and infectious virus in the WT mice compared to the PBS control treated WT mice (**Fig. 5c**). This trend was also observed when the FcRγ^-/-^ monocytes were transferred back into the FcRγ^-/-^ mice (**Fig. 5c**). In contrast, transfer of WT monocytes (CD45.1) into MAYV-infected WT or FcRγ^-/-^ CD45.2 recipient mice showed no change in infectious virus or viral RNA between monocyte transfer and PBS control for each genotype. Taken together, these data show that monocytes lacking activating FcγRs are sufficient to increase MAYV burden in joint-associated tissue indicating a pro-viral role for these cells.

**Figure 5:**
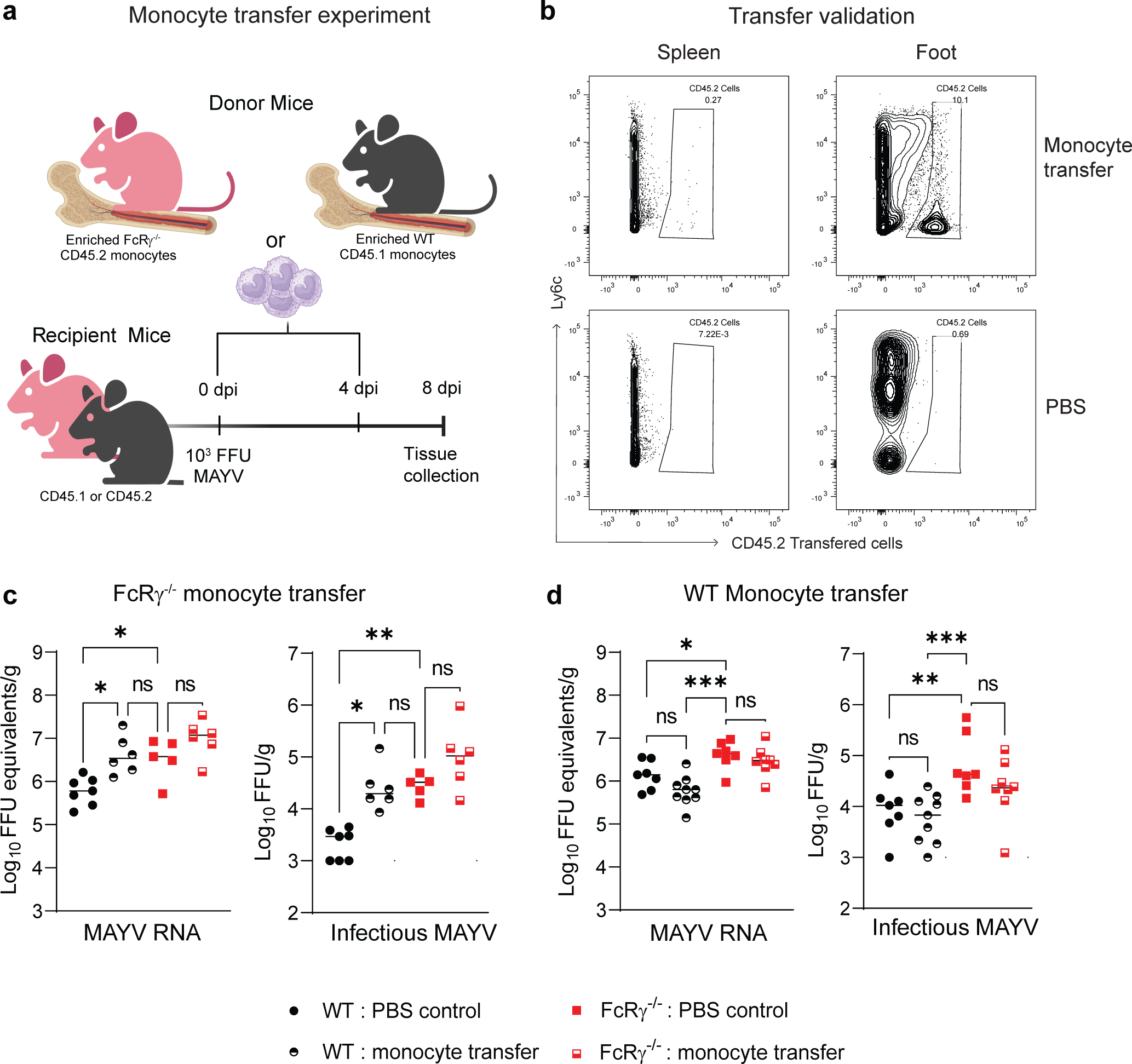
Monocytes lacking activating FcγRs enhance viral infection. (**a**) Schematic of monocyte transfer experiment. Monocytes were enriched through negative selection from the bone marrow of either FcRγ^−/−^ CD45.2 or WT CD45.1 mice. Recipient WT and FcRγ^−/−^ mice were injected intravenous with a PBS control or either WT or FcRγ^−/−^ monocytes at 0 dpi (5 x 10^5^ monocytes) and 4 dpi (1 x 10^6^ monocytes). (**b**) Representative flow plots show CD45.2 donor cells identified in single cell suspensions from either the spleen or contralateral ankle of CD45.1 recipient mice. (**c-d**) Quantification at 8 dpi of MAYV RNA and infectious virus from the ipsilateral ankle of mice receiving (**c**) FcRγ^−/−^ or (**d**) WT monocytes compared to PBS control. Bars indicate median values (n = 5 to 9; 2 to 3 independent experiments; Kruskal-Wallis test *, *P* < 0.05; **, *P* < 0.01; ***, *P* < 0.001).

## Discussion

The generation of anti-viral antibodies during alphavirus infection is critical for clearance of infectious virus; however, the importance of Fc effector functions to mediate this clearance was unknown. Here, we examined the role of activating FcγRs during MAYV infection. We determined that activating FcγRs are necessary for optimal resolution of clinical disease and clearance of MAYV RNA through 28 dpi, particularly for joint-associated tissues, which was not observed previously for CHIKV [27]. Analysis of antibody responses showed that despite equivalent neutralization and quantity at this chronic time point, significant increase in MAYV RNA remained in the tissue. Despite early differences in neutralizing antibodies, we showed that neutralizing antibodies were not sufficient to enhance viral clearance in FcRγ^−/−^ mice compared to mice that completely lack antibodies. Differences in monocyte infiltration as well as distinct genetic signatures were observed in FcRγ^−/−^ mice, and, when transferred in WT mice, the FcRγ^−/−^ monocytes were sufficient to increase levels of both infectious virus and viral RNA at this resolution time point. Taken together, these data demonstrate the necessity of FcγRs, specifically on monocytes, for protection during MAYV infection.

Fc effector functions have been shown to be important for protection from a variety of viruses [22, 70, 71]. In mice, NK cells, neutrophils, DCs, and monocytes express combinations of activating FcγRs (FcγRI, FcγRIII, and FcγRIV) that facilitate opsonization, antibody-dependent cellular cytotoxicity (ADCC), antibody-dependent cellular phagocytosis (ADCP), and/or enhance T cell activation through DC maturation through interaction with Fc region of IgG [72]. Previously, engagement of FcγRs on monocytes with antibodies was shown to be critical for mAb-dependent clearance of CHIKV and MAYV [19, 22, 29, 73]. Furthermore, non-neutralizing anti-MAYV mAbs increased phagocytosis of immune complexes into FcγR-bearing myeloid cells resulting in an abortive infection and clearance of the virus [30]. In our model, mice lacking the activating FcγRs showed prolonged disease and reduced MAYV clearance through chronic time points even in the presence of neutralizing antibodies. Interestingly, Jh mice, which fail to produce antibodies, had equivalent viral burden as the FcRγ^-/-^ mice in joint-associated tissues highlighting the host reliance on Fc-FcγR interactions for MAYV clearance rather than neutralizing antibodies for tissue-specific clearance. More broadly, studies have shown Fc-FcγR interaction on monocytes has also been implicated for enhanced mAb-mediated clearance during SARS-CoV-2 infection [74, 75], suggesting an importance to study these interactions.

Monocytes, neutrophils, and NK cells were increased at 8 and 10 dpi in FcRγ^−/−^ mice. This may be related to a delay in viral clearance in the absence Fc effector functions resulting in continued recruitment of the cells to the site of infection, although chemokine levels were similar between the groups (data not shown). Of particular interest was the inversion in the monocyte to macrophage ratio. By 10 dpi, WT mice had reduced numbers of monocytes with increased number of macrophages compared to FcRγ^−/−^ mice. Macrophages, which can be monocyte-derived, respond rapidly to stimuli from the environment and enact a diverse range of effector functions. Monocytes and macrophages have been shown to provide protective and pathogenic roles during alphavirus infection. Monocytes produce type I interferons following alphavirus recognition, which has been shown as necessary for controlling viral replication [73, 76, 77]. Depletion of phagocytes through clodronate-loaded liposomes prior to CHIKV infection reduced clinical disease [37]. A positive correlation between monocyte chemoattractant protein-1 (MCP-1; CCL2) levels and severe disease following CHIKV outbreaks has been recorded [78]. Indeed, inhibition of MCP-1 reduced CHIKV disease in mice [79]. Conversely, CCR2^-/-^ mice or depletion of CCR2^+^ inflammatory monocytes resulted in increased disease during MAYV, RRV, or CHIKV infections due to a compensatory influx of neutrophils and the potential for monocytes to still traffic into tissues through CCR7 [69, 77, 80, 81]. Our data suggests that FcγR engagement may limit the recruitment of monocytes/neutrophils promoting early disease resolution.

Despite lacking activating FcγRs in our model, the inhibitory FcγR (FcγRIIb) is still present since it does not signal through the Fc common gamma chain and it can be expressed on myeloid cells [33]. Interaction of the anti-viral antibodies with FcγRIIb, in the absence of the activating FcγRs, could have resulted in a differential cellular response. Previous work has shown an enhanced inflammatory response in the absence of FcγRIIb signaling [82]; however, the impact of only expressing FcγRIIb during infection is less understood. Taken together, the pathogenic or protection functions of monocytes may result from a more nuanced relationship between anti-viral antibodies and their interaction with the activating FcγRs.

Monocytes and macrophages have been identified as targets of alphavirus infection [40, 83–85] and specifically for MAYV [41, 86]. Interestingly, a recent study showed reduced MAYV burden in CCR2^-/-^ mice [81]. While it was hypothesized that the reduced viral load in the CCR2^-/-^ mice was related to an increase in neutrophil recruitment, it is possible that the recruited monocytes are an important source of infectious virus. Additionally, infiltrating monocytes are productive targets of CHIKV infection and increased infection in non-immune cells at the site of infection [85]. Notwithstanding, monocytes and macrophages can also phagocytose virally infected cells, so the presence of viral RNA does not necessarily equate to a productive infection. Since phagocytosis will most likely be reduced in the absence of activating FcγRs, it is unlikely that phagocytosis can fully explain the increase in viral RNA observed in the scRNA sequencing results. Pathway analysis showed enrichment of pathways utilized by alphaviruses for efficient viral replication, including proteasome-ubiquitination, oxidation, and heat stress response pathways. Indeed, treatment of cells with an oxidant during Sindbis virus infection enhanced viral RNA capping [87]. The nsP2 of CHIKV and other arthritogenic alphaviruses degrade the catalytic subunit of the RNA polymerase II through ubiquitination to block host transcription [88]. Inhibitors of ubiquitination or the heat shock response protein, HSP-90, reduced MAYV, Venezuelan equine encephalitis virus, and/or CHIKV replication [89–91]. Additionally, pathways associated with type I IFN, and anti-viral responses were enriched. Taken together, the pathway analysis results and the increased infectious MAYV following transfer of FcRγ^-/-^ monocytes suggest that lack of activating FcγR signaling alters the susceptibility of monocytes and macrophage to MAYV infection. Future studies are needed to identify specific genes and signaling pathways that mediate this enhancement infection which may provide insight into factors that promote susceptibility of myeloid cells to infections and ultimately how Fc-FcγR interactions modify cellular responses.

## Methods

### Cells and viruses

Vero cells (African Green Monkey Kidney, female; ATCC) were cultured in Dulbecco’s Modified Eagle Medium (DMEM) (Gibco) supplemented with 5% heat-inactivated fetal bovine serum (HI-FBS; Omega) at 37°C with 5% CO_2_. Mayaro virus isolate BeH407 was received from the World Reference Center for Emerging Viruses and Arboviruses (WRCEVA) and passaged twice on Vero cells. Ross River virus strain T48 was produced from an infectious cDNA clone, as described previously, and passaged once on BHK cells [92]. Chikungunya virus strain AF15561 was produced from an infectious cDNA clone, as previously described, and passaged once on BHK cells [93].

### Mouse Studies

All animal experiments and procedures were carried out in accordance with the recommendations in the Guide for the Care and Use of Laboratory Animals of the National Institutes of Health and approved by the NIAID ACUC under the protocol LVD 6E. Four-week-old male and female C57BL/6NTac (WT), B6.129P2-*Fcer1g^tm1Rav^*N12 (FcRγ^-/-^), C57BL/6NTac-*Igh-J^em1Tac^* (Jh), and B6.SJL-*Ptprc^a^*/BoyAiTac (Taconic Biosciences) were used for our studies. Footpad inoculations were performed under anesthesia that was induced and maintained with isoflurane. Retro-orbital intravenous injections were performed under anesthesia with 2, 2, 2-tribromoethanol (Avertin). Mice were inoculated subcutaneously in the rear footpad with 10^3^ FFU of MAYV, RRV, or CHIKV diluted in Hanks’ Balanced Salt Solution (HBSS, Gibco) supplemented with 1% HI-FBS. Foot swelling was measured (width x height) prior to infection and on indicated time points using digital calipers. Mice were sacrificed, perfused with PBS, and tissues collected at 3, 8, 10, or 28 dpi.

### Focus forming assay (FFA)

Tissues were weighed then homogenized in DMEM supplemented with 2% HI-FBS, 10mM HEPES (Gibco) and penicillin and streptomycin (Gibco) using a silica beads. Homogenates were clarified (12,000 × rpm for 5 min). Vero cells, plated in 96-well flat bottom plate one day prior, were infected with serial dilutions of clarified tissue homogenates for 2 hours at 37°C. The inoculum was removed, then the cells were overlayed with a 1% methylcellulose (Sigma-Aldrich) in Minimum Essential Medium (MEM, Sigma-Aldrich) supplemented with penicillin and streptomycin, 10 mM HEPES, and 2% HI-FBS. Cells were fixed 18 h later with 4% paraformaldehyde (PFA; Electron Microscopy Sciences) in PBS. Cells were washed with PBS, permeabilized with perm wash (PBS supplemented with 0.1% saponin and 0.1% BSA) and stained using CHK-48 [18]. Following a wash with ELISA wash buffer (PBS with 0.05% Tween-20), cells were incubated with peroxidase-conjugated goat anti-mouse IgG (H + L) antibody (SeraCare) for 1-2 h. Cells were washed with ELISA wash buffer and foci were developed using TrueBlue substrate (KPL) and counted using a Biospot plate reader (Cellular Technology, Inc.).

### Quantification of viral RNA

RNA was isolated using the KingFisher Duo Prime System with the MagMAX Viral RNA Isolation Kit (Applied Biosystems) following the manufacturer’s instructions. Viral RNA was quantified by qRT-PCR using the TaqMan Fast Virus 1-Step MasterMix with RRV nsp3 specific primers (Forward: 5’ - GTG TTC TCC GGA GGT AAA GAT AG -3’, Reverse: 5’ - TCG CGG CAA TAG ATG ACT AC - 3’) and probe (5’ - 6FAM/ACC TGT TTA/ZEN/CCG CAA TGG ACA CCA/ 3IABkFQ/ - 3’), CHIKV E1 specific primers (Forward: 5’ – TCG ACG CGC CAT CTT TAA – 3’, Reverse: 5’ – ATC GAA TGC ACC GCA CAC T – 3’) and probe (5’ – 6FAM/GCC GAG AGC/ZEN/CCG TTT TTA AAA TCA C/3IABkFQ – 3’) or MAYV E2 specific primers and probe (Forward: 5’ – GTG GTC GCA CAG TGA ATC TTT C- 3’, Reverse: 5’ – CAA ATG TCC ACC AGG CGA AG – 3’, Probe: 5’ - 6FAM /ATG GTG GTA/ZEN/GGC TAT CCC ACA GGT C/3IABkFQ – 3’) and compared to RNA isolated from viral stocks as a standard curve to determine FFU equivalents. Viral RNA was normalized to tissue weight.

### Focus reduction neutralization test (FRNT)

Serum from infected animals was serially diluted in DMEM supplemented with 2% HI-FBS, penicillin and streptomycin, and 10 mM HEPES and incubated with 10^2^ FFU MAYV for 1 h at 37°C in duplicate wells. Serum-virus mixtures were added to Vero cells for 90 min at 37°C followed by an overlay with a 1% methylcellulose in Minimum Essential Medium (MEM, Invitrogen) supplemented with penicillin and streptomycin, 10 mM HEPES, and 2% HI-FBS. Cells were fixed 18 h later after the addition of 4% PFA in PBS. Infected cells were incubated with CHK-48 (0.5 µg/ml). After washing and incubation with peroxidase-conjugated goat anti-mouse IgG (SeraCare), foci of infection were developed using TrueBlue substrate (KPL) and counted using a Biospot plate reader (Cellular Technology, Inc.). Wells containing serum dilutions were compared to wells inoculated in the absence of serum. The half maximal inhibitory serum dilution (Neut_50_ value) was calculated using non-linear regression analysis constraining the bottom to 0 and top to 100.

### Quantification of anti-MAYV E2 IgG in serum

Recombinant MAYV E2 protein (Native Antigen) was absorbed overnight at 4°C on Maxisorp immunocapture ELISA plates (Thermo Scientific) in a sodium bicarbonate buffer pH 9.3. Wells were washed with ELISA wash buffer and blocked with blocking buffer [PBS + 2% BSA (Sigma)] for 2 h at 37°C. Mouse serum was heat inactivated at 56°C for 1 h, serially diluted in blocking buffer, and then added to wells for 2 h at room temperature (RT). Plates were washed with ELISA wash buffer then incubated for 1 h at RT with an HRP conjugated goat anti-mouse (Southern Biotech). Plates were washed with ELISA wash buffer and developed using 1-Step Ultra TMB-ELISA substrate solution (Thermo Fisher). The reaction was stopped with 1 M H_2_SO_4_ and absorbance was measured at 450 nm. The EC_50_ of each sample was calculated using non-linear regression analysis after constraining the bottom to 0 and the top to 100 in GraphPad Prism.

### Antibody depletion of B cells

Mice were administered two doses of 500 μg mouse anti-mouse CD20 (BioXcell, clone MB20-11; cat# BE0356) or an isotype control (BioXcell, IgG2c) at 0 and 4 days post-infection by intraperitoneal injection. At 8 dpi, the ipsilateral and contralateral ankles were collected, homogenized, and viral burden was assessed by FFA as described above. Cells from peripheral blood were isolated and washed with FACS buffer (PBS with 1% BSA) and single cell suspensions were blocked for FcγR binding (BioLegend clone 93; 1:50) and then stained with the following antibodies to validate B cell depletion: CD45 BUV395 (BD Biosciences clone 30-F11; 1:200), CD19 BV737 (BD clone 1D3; 1:300), and B220/CD45R BV421 (Biolegend clone RA3-6B2; 1:200). Viability was determined through exclusion of a fixable viability dye (aqua). Samples were processed on a BD LSRFortessa (BD Biosciences) and analyzed using FlowJo (FlowJo, LLC).

### Serum IgG binding to live MAYV-infected cell surface

Vero cells were seeded at 2.5 x 10^4^ cells/well in 96 well plates and infected with MOI 5 MAYV for 18 h at 37°C with 5% CO_2._ Cells were then trypsinized until detached and resuspended in FACS buffer (PBS with 1% BSA). Cells were then incubated with serum diluted in FACS buffer (1:200) for 1 hr at 4°C. Cells were washed 3 times in FACS buffer and then stained with an AF647-conjugated anti-mouse IgG secondary antibody (eBiosciences; 1:2000). Cells were washed 3 times in FACS buffer then fixed in 4% PFA for 10 minutes at 4°C. Cells were then washed and run on a BD LSRFortessa.

### Flow Cytometry

Mice were infected with 10^3^ FFU of MAYV and at 3, 8, or 10 dpi mice were sacrificed and perfused with PBS. The ipsilateral feet were disarticulated, and the skin was everted. The skin was minced then the tissue (feet and skin combined) was digested in RPMI 1640 supplemented with collagenase (2.5mg/mL; Sigma), Liberase (100ug/mL; Roche), HEPES (15mM), and DNase I (10ug/mL; Sigma) for 2 h at 37°C with agitation, strained through a 70-μm filter, and resuspended in RPMI 1640 supplemented with 10% HI-FBS. Cells were then washed with FACS buffer, single cell suspensions were blocked for FcγR binding (BioLegend clone 93; 1:50), then stained with the following anti-mouse antibodies: CD45 BUV395 (BD Biosciences clone 30-F11; 1:200), CD3 Alexa488 (Biolegend clone KT3.1.1; 1:400), CD4 Alexa647 (eBiosciences clone RM4-5; 1:400), CD8b Alexa700 (eBiosciences clone YTS156.7.7; 1:400), NK1.1 PE-Cy7 (BioLegend clone PK136; 1:200), CD11b PerCP-Cy5.5 (BioLegend clone M1/70; 1:200), CD19 BV737 (BD clone 1D3; 1:300), Ly6C BV650 (biolegend clone HK1.4; 1:400), Ly6G APC-Cy7 (Biolegend clone 1A8; 1:300), CD11c PEcy5 (Biolegend clone N418; 1:400), I-A/I-E (MHC class II) BV711 (Biolegend clone M5/114.15.2; 1:400), and F4/80 BV421 (BD clone T45-2342; 1:400). Viability was determined through exclusion of a fixable viability dye (aqua). Samples were processed on a BD LSRFortessa and analyzed using FlowJo.

### Adoptive transfer of monocytes

The bone marrow from tibias and femurs of donor C57BL/6N CD45.1 or FcRy^-/-^ CD45.2 mice was aspirated and collected in RPMI 1640 (Invitrogen) at 4°C. Monocytes from bone marrow were enriched by negative selection (Monocyte Isolation Kit BM, Miltenyi Biotec) following the manufacturer’s instructions and resuspended in sterile PBS (Gibco). Negatively enriched monocytes were intravenously infused into C57BL/6N WT or FcRγ^-/-^ CD45.2 recipient mice at 0 dpi (5 x 10^6^ cells) and 4 dpi (1 x 10^6^ cells). At 8 dpi, the ipsilateral ankles were collected, homogenized, and viral burden was assessed by FFA and qRT-PCR as described above. The spleen and contralateral feet were collected to confirm monocyte transfer. The feet were digested, as described above. The spleen was passed through a 70 µm filter then rinsed with RPMI 1640 supplemented with 10% HI-FBS. Cells were then washed with FACS buffer, single cell suspensions were blocked for FcγR binding (BioLegend clone 93; 1:50), then stained with the following anti-mouse antibodies: CD45.1 BUV395 (BD Biosciences clone A20; 1:200), CD45.2 FITC (BD Biosciences clone 104; 1:200), CD11b PerCP-Cy5.5 (BioLegend clone M1/70; 1:200), and Ly6C BV650 (BioLegend clone HK1.4; 1:400). Viability was determined through exclusion of a fixable viability dye (aqua). Samples were processed on a BD LSRFortessa and analyzed using FlowJo.

### Single-cell RNAseq preparation and analysis

Mice were infected with 10^3^ FFU of MAYV and, at 10 dpi, mice were sacrificed, perfused with PBS, and the ipsilateral feet were dissociated into a single cell suspension, as described above. Cells were stained with anti-CD45 BUV395 (BD Biosciences clone 30-F11; 1:200) and a viability dye. Viable, unfixed, CD45^+^ cells were sorted on a BD FACSAria into RPMI supplemented with 10% FBS. Sorted CD45^+^ immune cells were centrifuged and resuspended in 1X PBS with BSA 0.04% to achieve 1000 cell/µL concentration. For the preparation of the cDNA and sequencing library generation, we followed manufacturer instructions from the Chromium Next GEM Single Cell 5’ Reagent Kit v2 (Dual index) User Guide with one modification: primers targeting the non-structural and subgenomic RNA were spiked-in during the cDNA synthesis to capture viral RNA (Step 1.1). The virus-specific primer concentration added to each RT reaction was ∼15pmoles [94]. All the other reagents were added according to the protocol, except the nuclease-free water, which was reduced to accommodate the primer spike-in volume. The Illumina library quality, yield, and size distribution was assessed by TapeStation D1000 high sensitivity assay, and by Qubit High Sensitivity dsDNA kit. The molar concentrations of the libraries were determined and diluted for sequencing according to Illumina sequencing protocol. We aimed to sequence each library to achieve **>**50,000 reads per cell. After sequencing, the fastQ files were submitted to Cell Ranger version 7.0.0 ‘mkref’, ‘mkfastq’ and ‘count’ functions with a custom genome of *Mus musculus* that contain the viral genome as exons (refdata-gex-mm10-2020-A). The filtered output of ‘counts’ (barcodes.tsv, features.tsv and matrix.mtx) were used for subsequent analysis with Seurat. In Seurat v4, we performed pre-processing of the data (quality controls) and normalization using the SCTransform function for accounting for batch effects. Once all samples were processed, they were integrated into one large Seurat object that contain all the conditions (two replicates of infected WT CD45^+^ cells, two replicates of infected FcRγ^-/-^ CD45^+^ cells, and one replicate of each mouse genotype as naive controls). We found 18 clusters at a clustering resolution of 0.4. We generated all marker genes of the clusters to assign the cell types within our dataset using the FindAllMarkers function in Seurat. Immune cell subsets were classified by a combination of the top 5 upregulated genes in each cluster as well as hallmark genes for certain cell types based on the literature (**Extended Data Fig. 3c-f**) [43–67]. For subsequent analysis of the myeloid cells, we subset the corresponding clusters ("0", "1", "2", "3", "4", "8", "16") into a new Seurat object (18,000 cells in total). We proceeded with re-analysis of this subset, resulting in 9 clusters. Cluster 8, corresponding to B cells, was removed from this analysis myeloid cells only. As described above, we used FindAllMarkers function to generate all markers genes for characterization of the cell types and subtypes (**Extended Data Fig. 4a-c**). Reads mapping to the viral template were counted and reads per cell were computed in Seurat, for both the sub-genomic and full-length genome, or aggregated as total viral reads, as shown in **Fig. 4g**.

### Pathway enrichment and modeling of gene networks

For differentially expressed gene (DEG) analysis, we used the myeloid cell Seurat objects, creating an extra column in the metadata that contained the Seurat cluster and the condition these cells came from, then we generated a dataframe containing all possible pairwise comparisons between clusters and conditions. From those, we selected the pairwise comparisons between infected FcRγ^-/-^ to WT for all clusters that had at least a minimum of 50 cells for each condition using the FindMarkers function as described in the differential expression testing vignette in the Seurat documentation. DEGs for comparisons between infected FcRγ^-/-^ to WT clusters were imported into Qiagen Ingenuity Pathways Analysis (IPA) (Ingenuity Systems; Qiagen, Redwood City, CA, USA). The list was subjected to a core analysis (-0.58 ≥ Log2FC ≥ 0.58, adjusted P value < 0.05), with significant IPA canonical pathways (p value < 0.05) assigned a Z score based on predicted activation state. The graphical summary of the canonical pathways highlights predicted interactions between terms, as curated by IPA.

### Similarity heatmaps of GO terms

Gene ontology analysis was performed on differentially expressed genes (absolute log2-fold change > 1 and FDR-adjusted p-value < 0.1) between infected FcRγ^-/-^ and WT clusters using the clusterProfiler v4.10.1 R package (https://doi.org/10.1089%2Fomi.2011.0118). Significantly enriched gene ontology pathways (FDR-adjusted p-value < 0.001) were then summarized using the simplifyEnrichment v1.12.0 R package (https://doi.org/10.1016/j.gpb.2022.04.008) to generate similarity heatmaps, revealing distinct sets of gene ontology terms with consistent similarities within each set.

### Statistical analysis

Statistical significance was assigned with P values using GraphPad Prism 9 (La Jolla, CA). Specific statistical test utilized are described in the figure legend of the corresponding data.

## Acknowledgements

This work was supported by the Intramural Research Program of NIAID/NIH.

## Data Availability

The data will be made available on a public repository. Additional requests should be directed to the corresponding author.

---

**Extended Data Figure 1: Contralateral foot swelling and viral RNA in tissues of FcR**γ**^-/-^ mice following MAYV infection.** Four-week-old WT or FcRγ^−/−^ C57BL/6N mice were infected subcutaneous in the rear footpad with 10^3^ focus forming units (FFU) of MAYV. (**a**) Swelling of the contralateral foot was measured prior to infection and for 25 dpi (n = 8 per group, 2 independent experiments). Graphs show mean ± SEM. Statistical significance was determined based on area under the curve (AUC) analysis using student’s t-test (ns = not significant). (**b**) Indicated tissues were harvested at 3 and 8 dpi and titrated for viral RNA by qRT-PCR with MAYV-specific primers and probe (n = 3 to 12 per group; 2 to 3 independent experiments). Statistical significance was determined by a Mann-Whitney test (**, *P* < 0.01; ****, *P* < 0.0001; ns = not significant). Bars indicate the median value and dotted lines indicate the limit of detection for the assay.

**Extended Data Figure 2: Flow gating scheme and adaptive immune responses**. (**a**) Flow gating scheme for identification of immune cell subsets. (**b**-**c**) Four-week-old WT or FcRγ^−/−^ C57BL/6N mice were infected subcutaneous in the rear footpad with 10^3^ FFU of MAYV. Single cell suspensions were isolated from the ipsilateral foot and proximal skin at (**b**) 3, 8, and 10 dpi or (**c**) 28 dpi stained for immune cells (CD45^+^), CD4 T cells (CD3^+^CD4^+^), CD8 T cells (CD3^+^CD8^+^), and B cells (CD3^-^CD19^+^) and analyzed by flow cytometry to determine the total numbers of viable cells or percentage of CD45^+^ cells (n = 5 to 8 per group; 3 independent experiments). (**c**) The gray bar represents the range of total cells and percentage of CD45^+^ cells from WT and FcRγ^−/−^ naïve mice. Graphs show mean ± SEM. Statistical significance was determined using a Mann-Whitney test at individual time points test. *, *P* <0.05; **, *P* < 0.01; ***, *P* < 0.001; ns = not significant.

**Extended Data Figure 3: Classification of immune cell clusters by genetic signature.** (**a**) Dot plot of the top 5 most signficant genes in each cluster from integrated RNA sequencing data, indicating log_2_FC and proportion of cells expressing each gene. (**b**) Cell identification of clusters based on additional key genes. (**c**) Dot plot of additional key gene identifiers in (b) showing log_2_FC and proportion of cells expressing each gene. (**d**) Distribution of cells across each cluster, shown for each individual mouse, indicating the log_2_(enrichment) of the clusters between the groups [n = 2 per infected condition, n = 1 for WT naive control, and n = 1 for FcRγ^-/-^ (KO) naive control]. Enrichment of B cells (Cluster 6) in the FcRγ^-/-^ naive sample is believed to be caused by a microbreak during intial tissue harvest, which is not present in any of the other samples

**Extended Data Figure 4: Classification of Myeloid subclusters by genetic signature.** (**a**) Dot plot of the top 5 most significant genes in the myeloid subcluster analysis, indicating log_2_FC and proportion of cells expressing each gene. (**b**) Additional key genes for cell identification of myeloid subcluster based on expert curation. (**c**) Dot plot of additional key gene identifiers in (b) showing log_2_FC and proportion of cells expressing each gene. (**d**) Distribution of cells across the subclusters, shown for each individual mouse, indicating the log_2_(enrichment) of the subclusters between the groups (n = 2 per infected condition, n = 1 for WT naive control, and n = 1 for FcRγ^-/-^ (KO) naive control). (**e**) The proportion of cells for each subcluster separated by genotype and the presence of MAYV RNA showing the log_2_(enrichment) of each subcluster between viral RNA positive and negative cells. (n = 2 per infected condition, n = 1 for WT naive control, and n = 1 for FcRγ^-/-^ (KO) naive control).

**Extended Data Figure 5: Overview of enriched pathways in FcR**γ**^-/-^ subclusters**. Differentially expressed genes (DEGs) enriched in FcRγ^-/-^ mice for each subcluster were analyzed using GO term analysis. Significant ontology terms were clustered based on semantic simillarity of member gene sets using simplifyEnrichment and hand annotated based on biological theme.

