## Supplementary figures and images for "Activating FcγRs on monocytes are necessary for optimal Mayaro virus clearance"

### Extended Data Fig 1

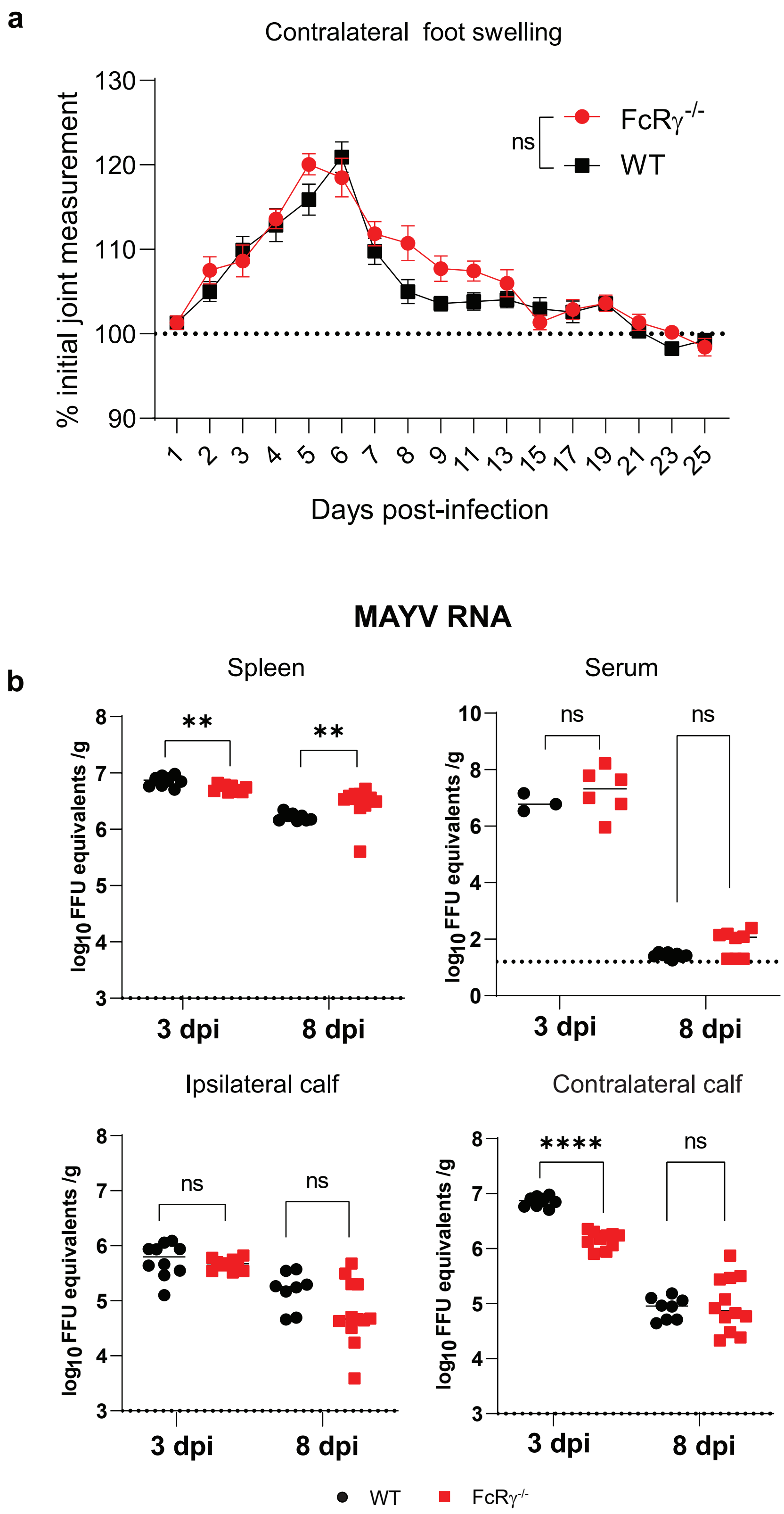


**Extended Data Figure 1**

### Extended Data Fig 2

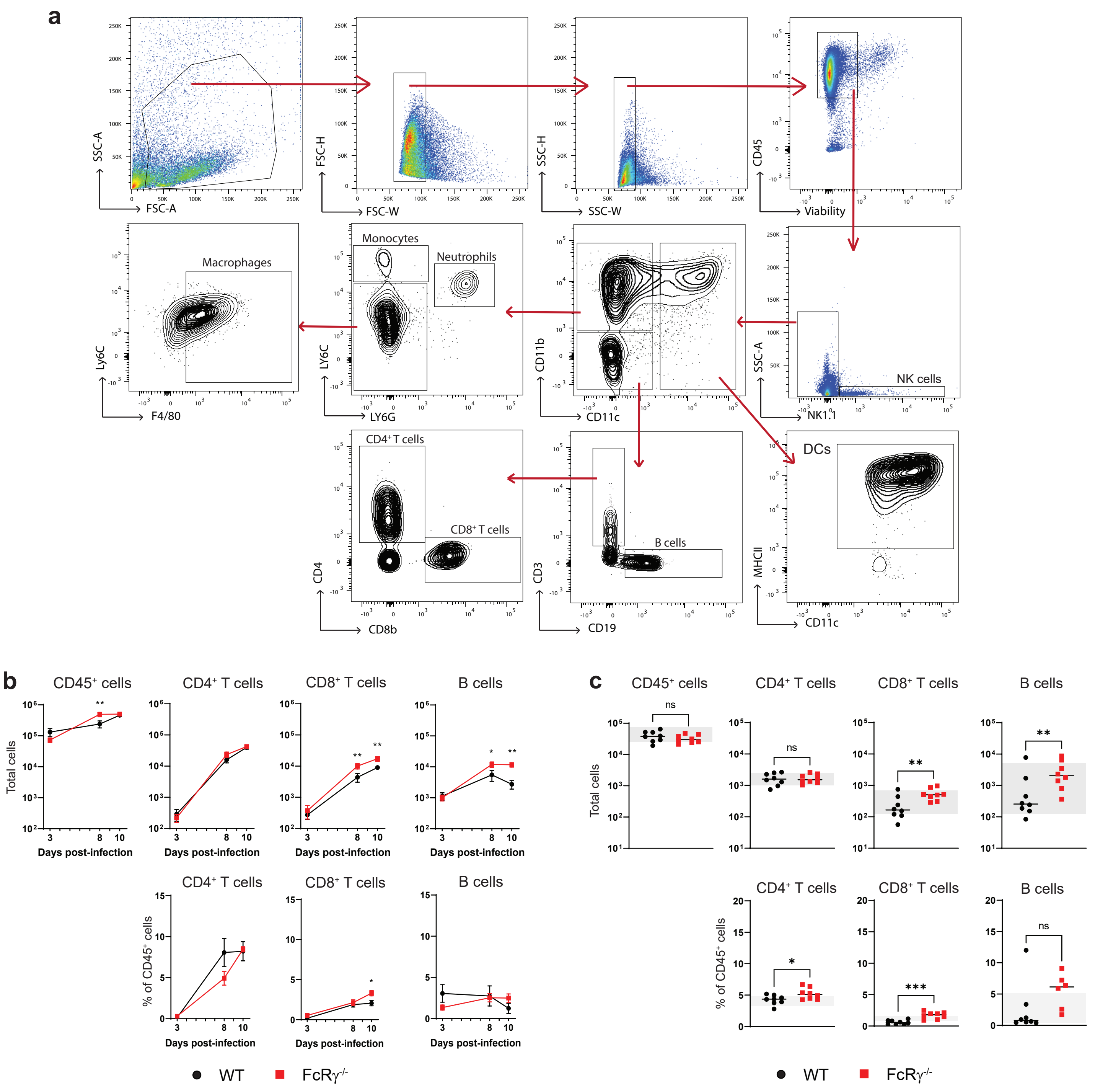


#### Extended Data Figure 2

### Extended Data Fig 5

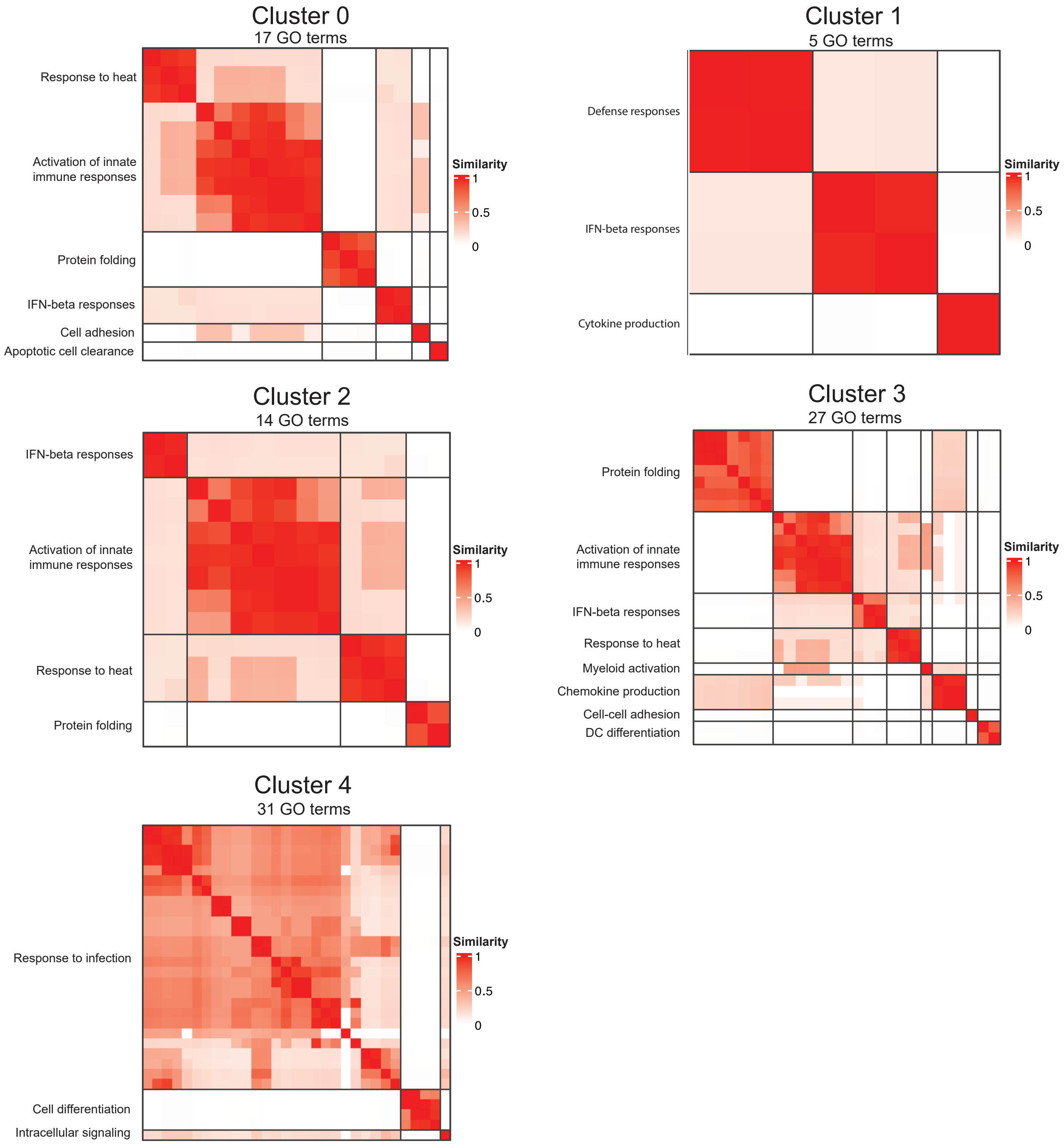


#### Extended Data Figure 5
