## Extended Data Fig 3 for "Activating FcγRs on monocytes are necessary for optimal Mayaro virus clearance"

**a** CD45⁺ cluster transcriptional markers

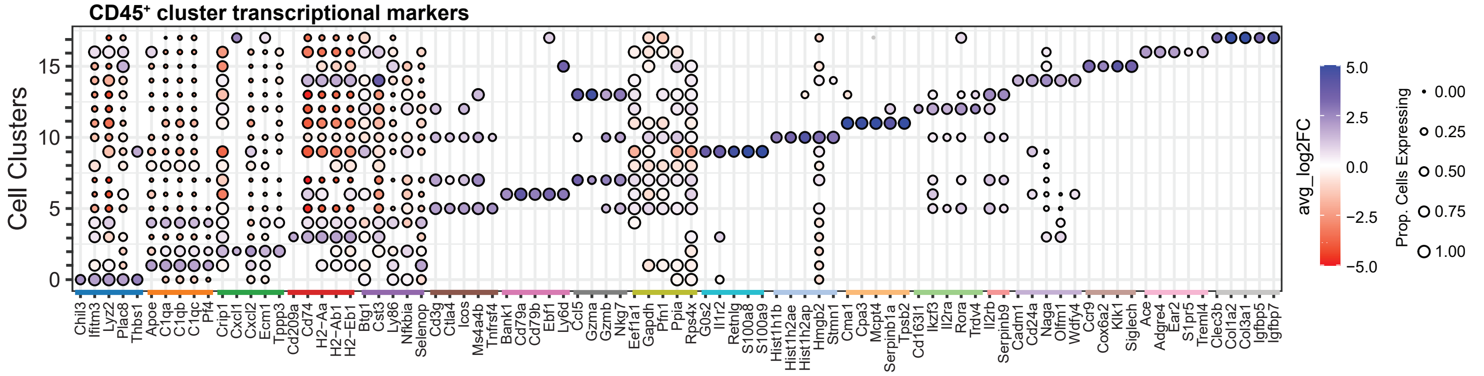


**b** Immune cell identification by cluster

| Cell type | Cluster # | Cell type | Additional key gene identifiers | References |
|---|---|---|---|---|
| **Dendritic Cells** | 3 | Resident DC | Flt3, Zbtb46, Cd209a, Itgax, H2-Aa | [43-46] |
| | 14 | Conventional DC | Flt3, Zbtb46, Xcr1, Itgax, Cadm1, Clec9a, H2-Aa | [44-46] |
| | 15 | Plasmacytoid DC | Siglech, Dntt, Pacsin1, Klk1, Itgax | [45-47] |
| **T cells** | 5 | CD4⁺ T cells | Cd4, Cd3d, Cd28, Ifng, Tnfrsf4, Gzmb | [45, 46, 48, 49, 67] |
| | 7 | CD8⁺ T cells | Cd8a, Cd8b1, CD3d, Cd28, Gzma, Gzmb | [45, 46, 48] |
| | 10 | T cells | Cd4, Cd8b1, Cd3d, Cd28, Ifng, Tnfrsf4, Gzmb | [45, 46, 48] |
| | 12 | γδ T cells | Tcrg-V6, Cd3d, Aqp3 | [45, 46, 50] |
| **Monocytes/ Macrophage** | 0 | Monocyte | Cd14, Chil3, Ly6c2, Csf1r, Ccr2 | [44-47] |
| | 1 | Macrophage | Cd14, Adgre1, Cd68, Mrc1, C1qb, H2-Aa, Itgam, Csf1r, Lgmn | [44-46, 51, 65, 66] |
| | 2 | Macrophage | Cd14, Cd68, C1qb, Itgam, Csf1r | [44-46] |
| | 4 | Macrophage | Adgre1, Cd68, Mrc1, C1qb, H2-Aa, Itgam, Ccr2, Csf1r, Lgmn | [44-46, 51] |
| | 8 | Macrophage | Lgmn | [45, 46, 52, 53] |
| | 16 | Non-classical monocyte | Itgam, Itgax, Ace, Ear2, Cx3cr1, Csf1r, Cd36 | [46, 54] |
| **B cells** | 6 | B cell | Cd19, Fcer2a, Cd79a | [46, 55] |
| **Neutrophils** | 9 | Neutrophil | Mmp9, Wfdc21, Siglece, Cxcr2, Itgam | [46, 55, 56] - |
| **Mast cells** | 11 | Mast cell | Mrgprb1, Kit | [46, 57, 58] |
| **NK cells** | 13 | NK cell | Ncr1, Klra8, Ifng, Itgax, Gzma, Gzmb | [46, 55, 59] |
| **Fibroblasts** | 17 | Fibroblast | Clec3b, Eln, Postn, Col1a1, Dcn | [46, 55] |

**c** Table Transcriptional Markers

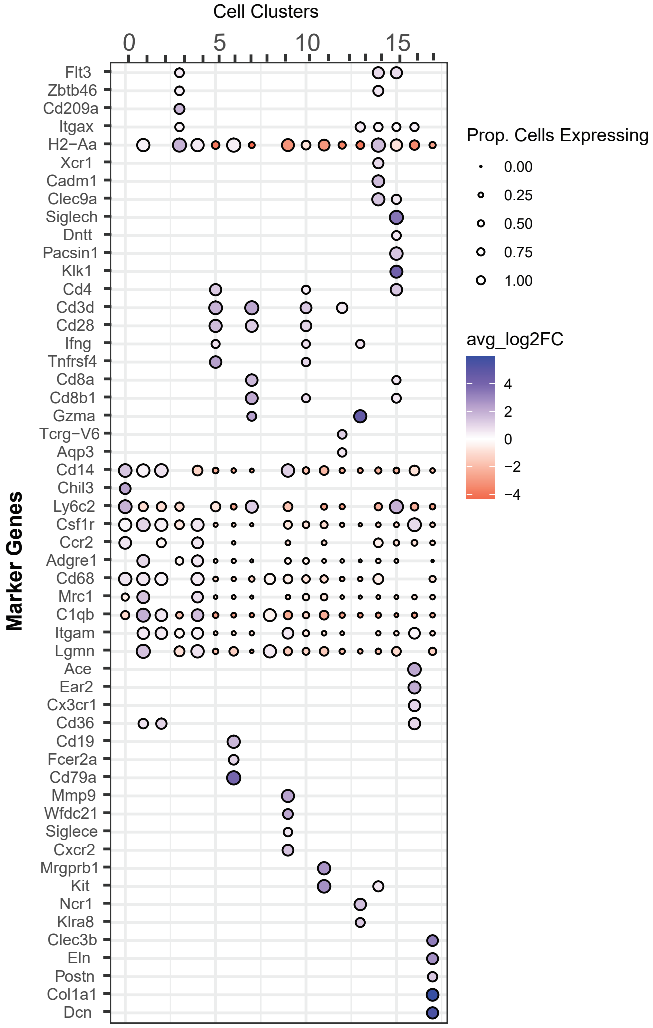


**d** Distribution of cells across clusters

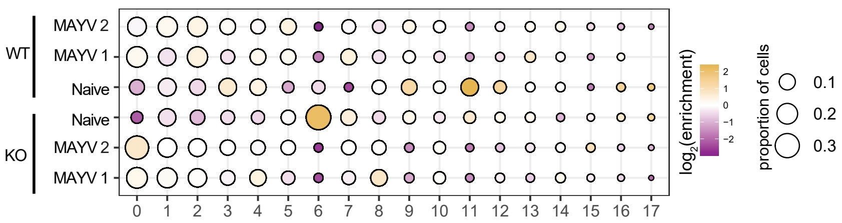


**Extended Data Figure 3**
