## Extended Data Fig 4 for "Activating FcγRs on monocytes are necessary for optimal Mayaro virus clearance"

**a** **Myeloid cell markers**

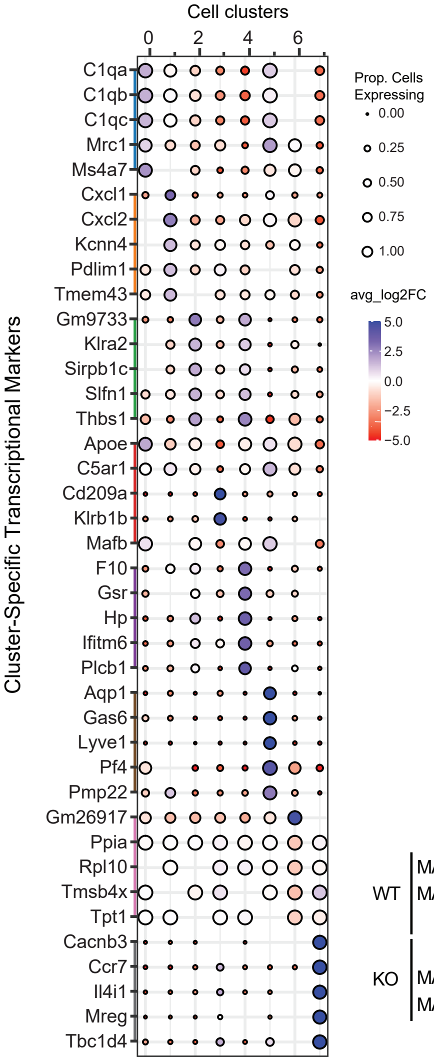


**b** **Immune cell identification by cluster**

| Cluster # | Cell type | Additional key gene identifiers | References |
|---|---|---|---|
| 0 | M2-like macrophage | C1qb, Adgre1, Trem2, Mrc1, Itgam, H2-Aa, Cd68, Cx3cr1, Il10ra | [44, 46, 51, 60, 66] |
| 1 | M1-like macrophage | Cxcl1, Cxcl2, Tnf, Trem2, Itgam, C1qb | [46, 60-62] |
| 2 | Inflammatory monocyte | Ly6i, Ly6c2, Chil3, Isg15, Ccr2, H2-Aa, Cd86, Cd68, Il10ra, Adgre1 | [44, 46, 47] |
| 3 | Activated DC | Cd209a, Zbtb46, Flt3, Itgax, H2-Aa, Ccr7, Cd163, Cd86 | [46, 60] |
| 4 | Non-classical monocyte | Ace, Chil3, Ly6c2, Ear2, Cd86, Isg15, Cd68, Il10ra, Ly6i, Ccr2, Cx3cr1, Cd86 | [44, 46, 47] |
| 5 | Tissue-resident macrophage | Lyve1, Cd163, Mrc1, C1qb, Trem2, Tnf | [46, 51, 60, 66] |
| 6 | Macrophage | Mrc1, Itgad | [44, 46] |
| 7 | DC | Ccr7, Fscn1, Il12b, Apol7c, Flt3, Zbtb46, H2-Aa, Apol7c | [46, 60, 63-65] |

**c** **Table Transcriptional Markers**

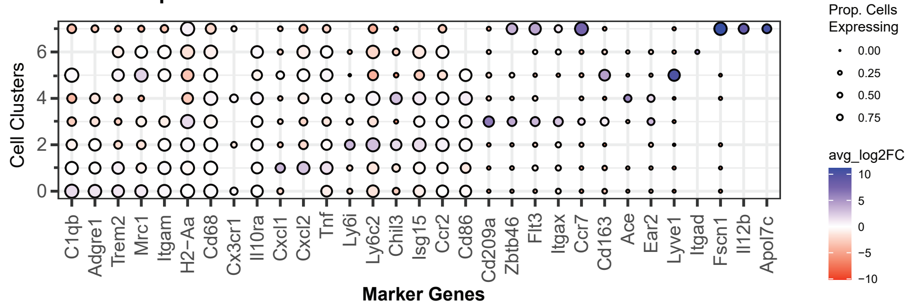


**d** **Distribution of cells across myeloid subclusters**

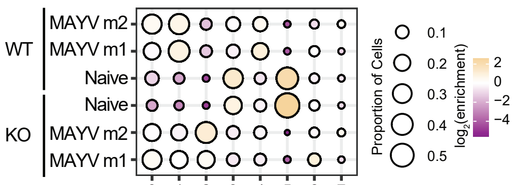


**e** **Distribution of MAYV⁺ cells across myeloid subclusters**

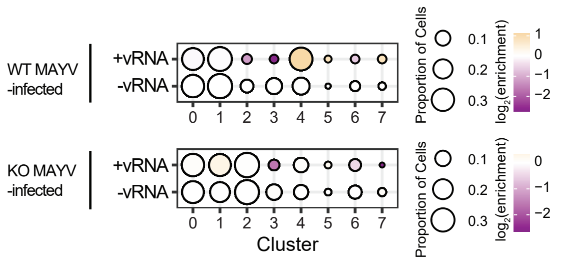


### Extended Data Figure 4
